# Intracranial interictal functional networks predict the effect of subacute cortical stimulation in focal epilepsy

**DOI:** 10.1101/288019

**Authors:** Yujiang Wang, Peter Neal Taylor, Judith Díaz-Díaz, Richard P. Selway, Antonio Valentin, Gonzalo Alarcón, Diego Jiménez-Jiménez

## Abstract

**Background:** Electric neurostimulation is being developed as an alternative treatment for drug-resistant epilepsy patients. A major challenge is to identify, among all possible combinations of stimulation parameters, the most effective stimulation protocol to achieve seizure control for each patient.

**Objective:** To estimate the value of interictal intracranial EEG recordings in identifying the most effective stimulation protocol in order to decrease number of seizures during intracranial subacute cortical stimulation (SCS).

**Methods:** Five patients undergoing SCS were included in this retrospective study. Bivariate correlation measures were applied to baseline interictal intracranial EEG recordings to infer functional correlation networks. The node strength of these networks was calculated for each recording electrode (indicating the overall correlation between activity from one electrode with the activity from all other electrodes). The relationship between node strength and the change in seizure rate associated with SCS was studied for the electrodes used for stimulation.

**Results:** Across all five patients, node strength was negatively correlated with seizure rate ratio (ratio of seizure rate during SCS relative to during baseline) for all frequency bands tested (delta, theta, alpha, beta, gamma, and high gamma), and both the common reference and the common average montage.

**Conclusion:** Interictal functional correlation networks contain information that correlates with the effect of SCS on the rate of epileptic seizures, suggesting that stimulation of those regions with higher node strength (derived from correlation networks) is more effective in reducing number of seizures.

**HIGHLIGHTS:**

- In focal epilepsy, effective brain stimulation targets are lacking
- We use baseline interictal functional networks to outline stimulation targets
- Stimulation of regions with higher node strength is most effective in our study

## Introduction

In approximately 35% of patients with epilepsy, seizures are not satisfactorily controlled by antiepileptic drugs [1, 2]. In the drug resistant group, resective epilepsy surgery is one treatment option [3]. However, in some patients this is not possible, e.g. due to the overlap of the hypothesised epileptogenic zone with eloquent cortex, which when removed would severely impair functional abilities. For such patients, electrical neurostimulation can be an alternative treatment option to supress their seizures [4]. A variety of stimulation strategies have been proposed, with different stimulation targets, intensities, frequencies, etc.; however, stimulation success rate has varied between studies (e.g. [5–10]). Presently, our understanding of the mechanism of electric neurostimulation is limited in epilepsy [11], and there is no consensus on how to identify a successful neurostimulation protocol. One difficulty is the enormous number of parameter combinations available (in terms of stimulation location, intensity or duration [9]), especially as the successful stimulation parameters may be patient-specific.

One approach is to perform electrical stimulation during intracranial recordings (subacute cortical stimulation or SCS) in order to identify the best stimulation location and parameters which, if effective, could later be used for chronic treatment with electrical stimulation [9, 10, 12]. For SCS, intracranial electroencephalography (iEEG) is performed as part of the routine presurgical assessment. In case resective surgery is discouraged, a pair of electrodes are selected which delivers electric stimulation while all remaining electrodes continue recording ongoing brain activity. A recent retrospective study showed that SCS in frontal lobe epilepsy was effective in 4 out of 5 patients, with at least 50% reduction in number of seizures in at least one stimulation setting [9]. Recently, a retrospective study of chronic stimulation using very similar parameters reported successful suppression of interictal discharges in 13 patients, with 10 out of 13 patients also reporting improvement in epilepsy severity and life satisfaction [10].

Alongside these studies on invasive electric neurostimulation, recent studies have shown that interictal recordings may contain more information about the epileptogenic areas and networks than standard clinical assessment can reveal (e.g. [13–17]). From interictal recordings, these studies have derived functional networks which capture how activity at one location relates to that of another location. Based on these networks and associated computational models, the studies elucidated network properties of the epileptogenic region, and proposed techniques to better localise such regions. For example, in [16], the authors used interictal intracranial recordings to derive a functional network, based on which they simulated seizure likelihood of each electrode position after epilepsy surgery patient-specifically. The simulation results were predictive of the actual surgical outcome in these patients with 81% accuracy. Meanwhile, in [17], the authors used pre-ictal functional networks to demonstrate that “push-pull” network interactions can predict if the following seizure will secondarily generalise.

Combining these insights with the recent proposal by [18] that such network analysis techniques may be informative for identifying suitable neurostimulation targets, we propose to use the SCS dataset in a retrospective functional network analysis. Specifically, we will investigate if interictal functional networks at baseline (before stimulation) can predict the success of the stimulation location in suppressing seizures.

## Material and methods

### Patient EEG and SCS data

The study used retrospective data from patients who underwent invasive pre-surgical assessment for the treatment of epilepsy in the Department of Clinical Neurophysiology at King’s College Hospital, London and fulfilled the following inclusion criteria:

a. Patients who had intracranial electrodes,
b. Patients not candidates for surgical resection,
c. Patients who underwent Subacute Cortical Stimulation for longer than 24 hours.

Data from patients where SCS was shortened due to pain or discomfort were excluded from the study.

All the patients were fully informed of the nature of the research according to the Declaration of Helsinki. The experimental procedure was approved by the ethical committee of Health Research Authority (IRAS 187072).

### Electrode implantation and recording protocol

In accordance to the inclusion criteria above, all patients had either subdural or depth electrodes implanted for intracranial EEG recordings during pre-surgical assessment. The number of electrodes implanted was necessarily limited by clinical practice. The position of the electrodes was determined by the suspected location of the ictal onset region, according to non-invasive evaluation: clinical history, scalp EEG recordings, neuropsychology, and neuroimaging [19]. The implantation procedure has been described elsewhere [20].

Recording of iEEG started when the patient recovered from the general anaesthesia, at least 24 hours after the surgery for electrode implantation. Cable telemetry with up to 64 recording channels was used for data acquisition with simultaneous video monitoring. In all patients, a Medelec-Profile system was used (Medelec, Oxford Instruments, United Kingdom). Data were digitized at 1024 Hz and high pass filtered at 0.5 Hz. The input range was 10 mV and data were digitized with a 22-bit analog-to-digital converter (an amplitude resolution of 0.153 nV). Interictal awake and sleep recordings in addition to ictal recordings were permanently stored. Data were recorded as referenced to a scalp electrode applied half way between Cz and Pz.

### SCS protocol

After sufficient seizures were recorded for clinical purposes (presurgical assessment), and the clinical team decided against resective surgery (e.g. due to proximity of the hypothesised epileptogenic cortex to eloquent cortex), SCS was performed. SCS was usually using different conditions (i.e. combinations of stimulation parameters and cortical locations) separated by breaks of no stimulation (Fig. 1A). In each condition one pair of electrodes was selected to be stimulated.

**Figure 1:**
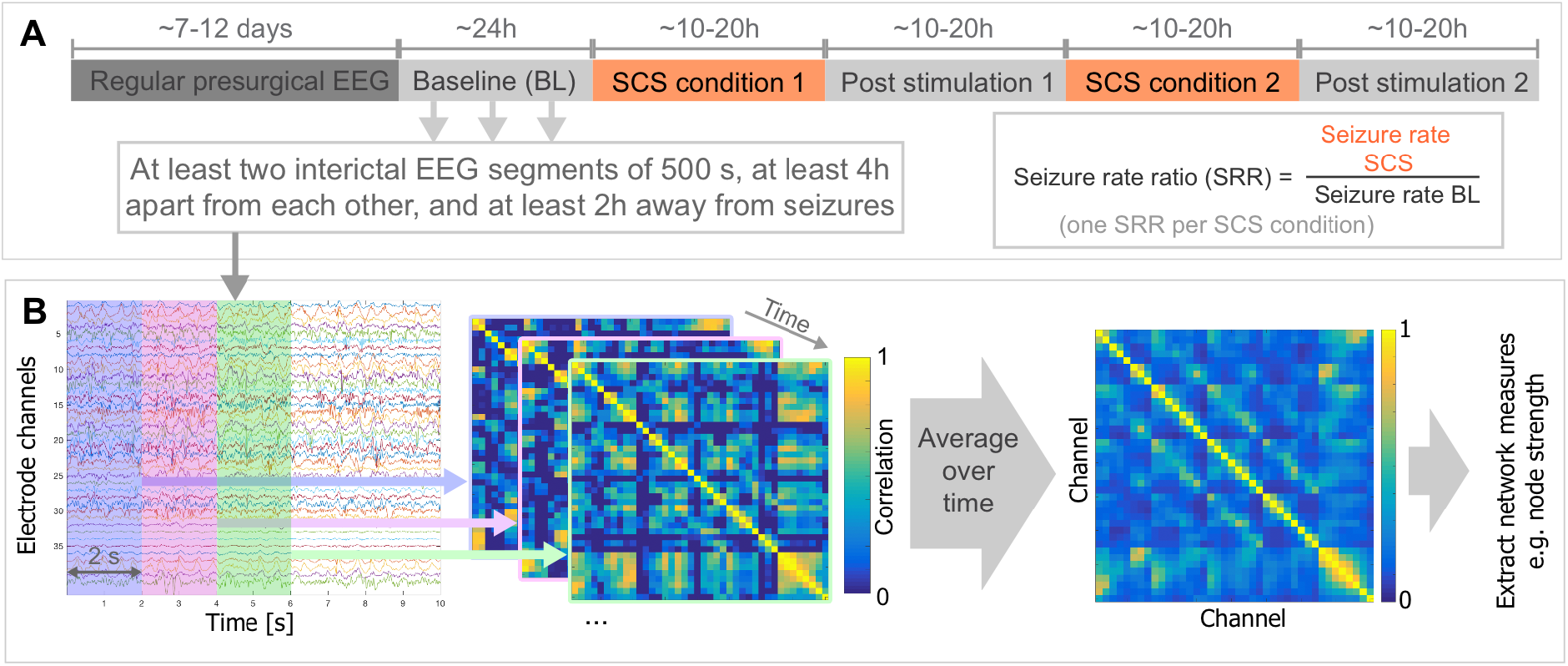
Schematic illustration of **A** experimental setup, and **B** data analysis pipeline.

Stimulation was carried out with a Medtronic stimulator set with the following parameters: frequency between 60 and 180 Hz, intensity between 0.5 and 3 mA, pulse duration between 90 and 450 µsec of biphasic continuous stimulation, usually for several hours. Cortical targets for SCS were selected independently of the presented analysis in this paper, and based on the following clinical criteria: a) abnormal responses to single pulse electrical stimulation (SPES) [21]; b) areas involved in electrographic seizure onset; c) areas showing the most frequent interictal discharges or areas close to interictal focal slow activity and d) areas showing an MRI lesion [9].

### Retrospective data analysis

For each patient, at least two interictal iEEG segments of 500 seconds were used from the baseline condition. These segments were also at least 4 hours apart from each other and 2 hours away from seizures (illustrated in Fig. 1A). Note that the baseline condition is essentially a period of about 24 hours after enough data has been collected for pre-surgical assessment purposes (but the clinical team has decided against resective surgery at this point), but before the first stimulation condition. In the baseline condition, the patients still continued to have seizures and were treated often with a combination of drugs (see Table 2 for details).

#### Seizure rate ratio

To determine the effect of SCS, seizure rates were counted (in number of seizures per hour) for baseline and for each stimulation condition by trained clinical team members (DJJ and JDD).

To determine the change in seizure rate from baseline to each SCS condition, we calculated the seizure rate ratio (SRR) as the ratio of the seizure rate in a SCS condition over the seizure rate during baseline. An SRR of 1 indicates no change in seizure rate. SRRs larger (smaller) than 1 indicates seizure rate increase (reduction) during the SCS condition.

#### Signal processing

As a first step, we visually inspected our data for noisy channels and excluded those channels from the analysis. Furthermore, intracranial depth electrodes located in white matter (determined by CT & MRI) were also excluded from analysis. To obtain a common average reference, we subtracted the mean of all channels from each channel. Frequency filtering was performed at the following frequency bands using fourth order Butterworth bandpass zero-phase digital filtering: delta 1-4 Hz, theta 4-8 Hz, alpha 8-13 Hz, beta 13-30 Hz, gamma 30-60 Hz, and high gamma 60-90 Hz.

For our main analysis, we focused on the alpha bandpass filtered data using a common average reference. Analyses were also carried out for all other frequency bands and for the common reference montage.

#### Functional (correlation) networks

To understand the spatio-temporal iEEG data from the baseline condition, we performed a functional network analysis (Fig. 1B). This is a method of analysing the bivariate properties of activity of channels over time (e.g. how correlated the activity is in one channel to another) as a matrix/network/graph [22]. Such functional network analyses approaches have been useful for analysing the spatial and temporal interactions of EEG data in the literature (see e.g. [23, 24, 16]). To create functional networks, we split up each 500 second segment of iEEG data into 2 second (non-overlapping) windows, and computed a Pearson correlation matrix for each window. These matrices were averaged over the entire 500 second segment to obtain a temporally averaged correlation network for each segment. It has been shown that a temporally invariant, or stable, network emerges after about 60-100 seconds [25]; hence we chose 500 seconds to ensure reliability (see also Appendix A for more details on reliability and temporal variations). We then removed the negative correlations from this matrix, and averaged all matrices across segments from the baseline condition in the same subject (following previous work [16]). This average matrix from the baseline recording is the functional network we will use for each subject for all subsequent analysis in the Results section.

As a network measure, we focused on node strength, which is the summed weights of all connections associated with a node. This choice is due to the interpretation of this network measure: it represents how strongly connected a node is to all other nodes in the network; in other words, how strongly correlated the activity in one channel is overall to all other channels’ activities

### Statistical analysis

To enable comparison across subjects, we standardize the node strength for each subject, so that node strength is transformed into a z-score. A positive (negative) value means that the node is more (less) connected than the average node in the network, and the value is in units of standard deviation.

To relate the network to seizure rate ratios, we computed the Pearson’s correlation between the average standardized node strength of the two stimulated channels in each condition and the seizure rate ratios of the corresponding condition across subjects. We obtained correlation coefficients for all frequency bands and for both the common reference and the common average montage.

## Results

### Patients

Our retrospective study included 5 patients, 1 male and 4 females with an average age of 12 years (range 7 - 22) who fulfilled the inclusion and exclusion criteria. Table 1 shows the patient characteristics. Table 2 shows the stimulation locations and parameters used for SCS.

**Table 1:** Summary of patient’s clinical details. SCS = Subacute Cortical Stimulation, M=male, F=female, F=Frontal lobe, T=Temporal lobe, P=Parietal lobe, FP=Frontal and parietal lobes, H=Hippocampus, STS=Superior temporal sulcus, MTG=Middle temporal gyrus, TOP=Temporal, occipital, and parietal lobes.

| Patient number | Gender | Age | Age of onset | Aetiology | Electrode setup |
| --- | --- | --- | --- | --- | --- |
| 1 | F | 7 years old | 4 years | Cortical Dysplasia | Six 8-channel depth shanks in the FP, and one 4-channel strip in F. |
| 2 | F | 15 years old | 11 years | Cortical Dysplasia | One 32-channel grid in FP, three 8-channel depth shanks in FP, and one 4-channel strip in the FP. |
| 3 | F | 9 years old | 8 months | Tuberous Sclerosis | One 32-channel grid in FP, three 8-channel strips in the FP, and one 4-channel strip in P. |
| 4 | M | 7 years old | 5 years | Cortical Dysplasia | One 8-channel strip in F, one 20-channel grid in F, one 4-channel strip in F, and one 4-channel strip in T. |
| 5 | M | 18 years old | 12 years | Unknown | Four 10-channel depth shanks in H, three 10-channel depth shanks in STS MTG, and four 10-channel depth shanks in TOP. |

**Table 2:**
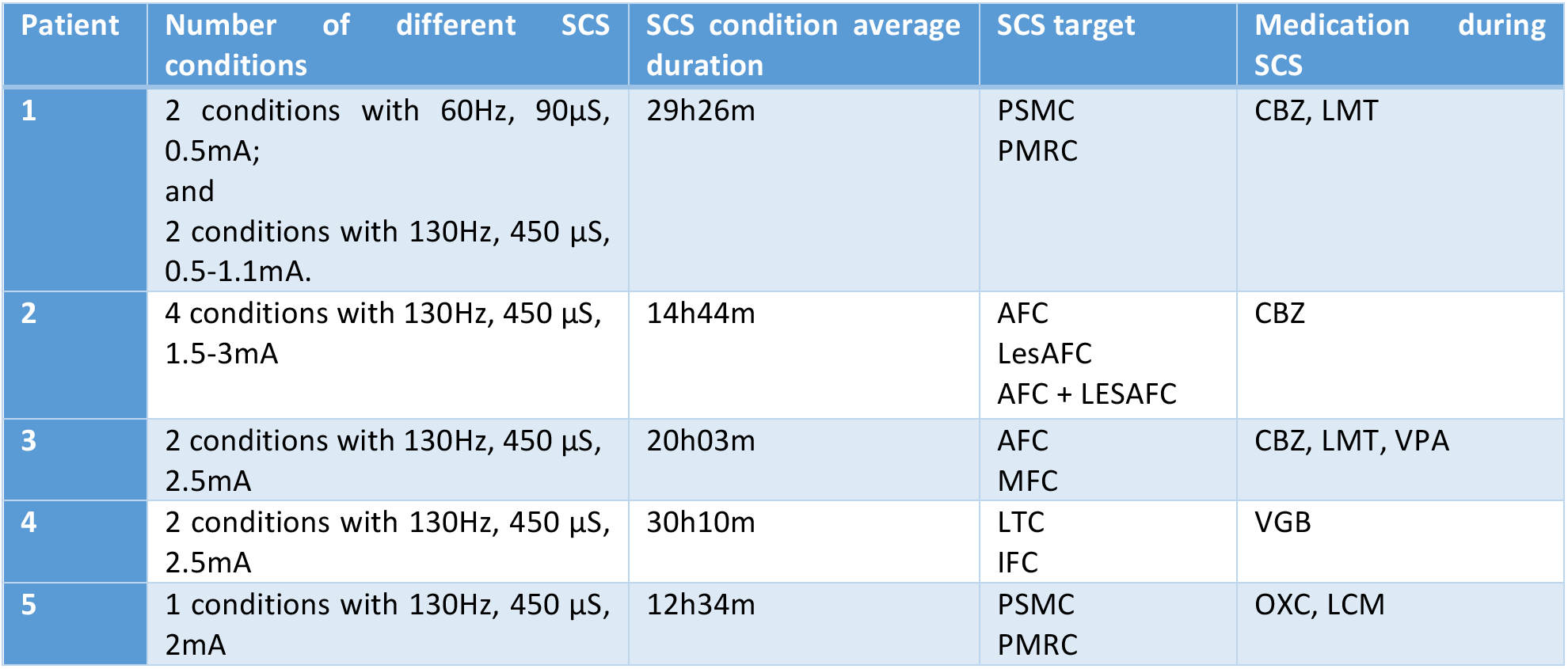
Summary of SCS condition average duration and parameters used in every patient. PSMC=Primary somatosensory cortex, PMRC=Primary motor cortex, AFC=Anterior frontal cortex, LesAFC=Lesional anterior frontal cortex, MFC=Middle frontal cortex, LTC=Lateral temporal cortex, IFC=Inferior frontal cortex, CBZ=Carbamazepine, LMT=Lamotrigine, VPA=Valproate, OXC=Oxcarbazepine, LSM=Lacosamide,

| Patient | Number of different SCS conditions | SCS condition average duration | SCS target | Medication during SCS |
| --- | --- | --- | --- | --- |
| 1 | 2 conditions with 60Hz, 90μS, 0.5mA; and 2 conditions with 130Hz, 450 μS, 0.5-1.1mA. | 29h26m | PSMC<br>PMRC | CBZ, LMT |
| 2 | 4 conditions with 130Hz, 450 μS, 1.5-3mA | 14h44m | AFC<br>LesAFC<br>AFC + LESAFC | CBZ |
| 3 | 2 conditions with 130Hz, 450 μS, 2.5mA | 20h03m | AFC<br>MFC | CBZ, LMT, VPA |
| 4 | 2 conditions with 130Hz, 450 μS, 2.5mA | 30h10m | LTC<br>IFC | VGB |
| 5 | 1 conditions with 130Hz, 450 μS, 2mA | 12h34m | PSMC<br>PMRC | OXC, LCM |

### Interictal network properties for one example subject

We begin our analysis by investigating the interictal baseline alpha band network properties for one example subject (Patient 2, see Table 1 and 2 for further details), who had a grid, a strip and three depth shanks implanted (Fig. 2A). We observe in this patient that the most anterior (grid and depth) electrodes show a high node strength (red in Fig. 2A). The activity in each of these electrodes are highly correlated to that of other electrodes on average. Conversely, the activity of the most posterior electrodes on the grid and also the strip electrodes in Fig. 2A show relatively (very) little correlation to other electrodes’ activities on average (coloured in (dark) blue Fig. 2A).

**Figure 2:**
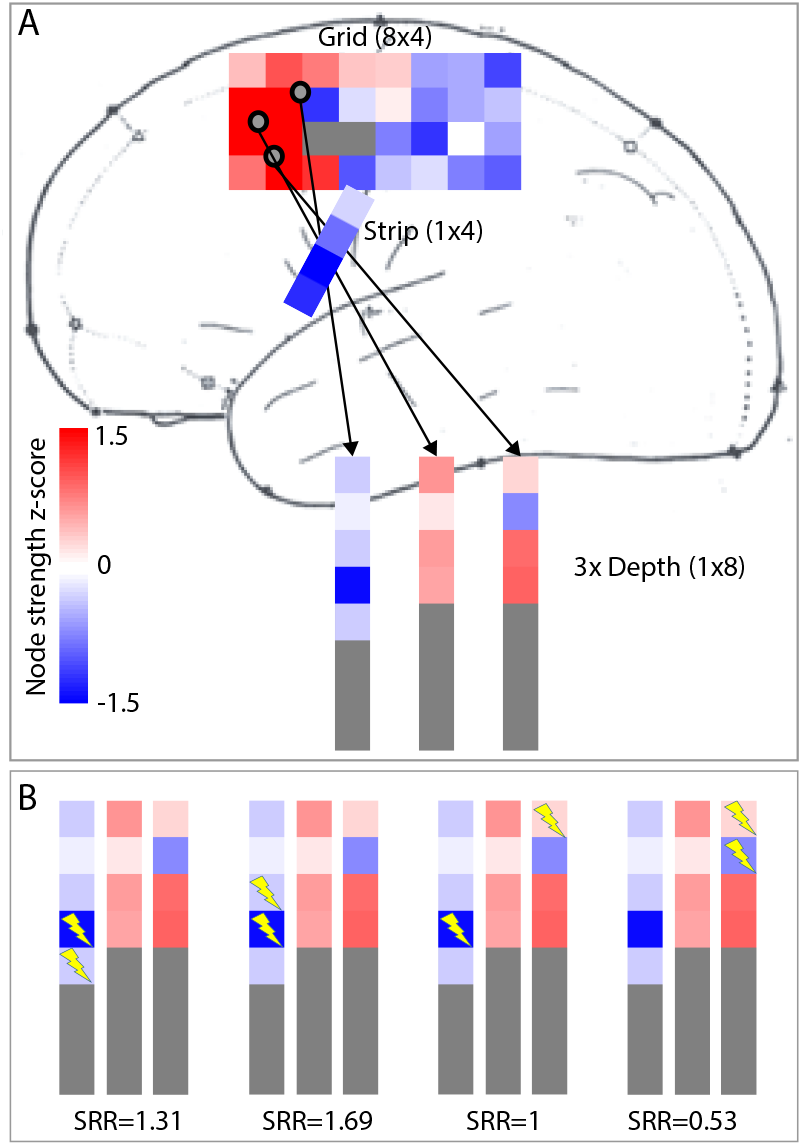
Interictal network property of one example subject in the alpha band. **A**: Node strength of the network shown on top of the schematic electrode layout, colour-coded according to the z-score. Grey indicates electrodes that have been excluded from the analysis due to noise, or because the electrode was located in the white matter. The three depth shanks are shown outside of the brain schematic for clarity. **B**: Four different stimulation conditions (all applied to electrodes on the depth shanks), and the stimulation electrode pairs are marked by yellow thunderbolt symbols. For simplicity, only the three depth shanks are shown, and the display is in the same order and orientation as in A.

To evaluate if there is a relationship between the interictal network node strength and the SCS locations, we overlaid the stimulation electrode pairs in each stimulation condition in this patient (Fig. 2B). Generally, when blue or dark blue electrodes (i.e. low node strength) are used for stimulation, the seizure rates increased during the stimulation (Fig. 2B first two panels). In one SCS (third panel in Fig. 2B), a dark blue and a light red electrode were used to stimulate. In this case, the seizure rate stayed the same as compared to baseline. Finally, when a light blue and a light red electrode were used to stimulate (fourth panel in Fig. 2B), a decrease of the seizure rate of almost half compared to baseline was observed.

### Interictal network node strength correlates with SCS effect

To perform our analysis across all five subjects, we correlated the z-score of interictal baseline alpha band network node strength of the two target electrodes in each SCS condition with the seizure rate ratio of the condition. Based on the example in the previous section, we wanted to test if there is a consistent correlation across patients and SCS conditions. Fig. 3A shows the scatterplot of z-score of node strength vs. seizure rate ratio for all patients and their different SCS conditions. We found a significant correlation between the two quantities (r=0.56). We also investigated the two striking outliers of two SCS conditions in patient 1 (outside of the confidence interval of the correlation, marked by yellow crosses in Fig. 3A). These two SCS conditions were performed at 60 Hz stimulation frequency, where all other SCS conditions shown were at 130 Hz. When excluding these two data points, the overall correlation coefficient increases to r=-0.711, with a p-value of 0.014.

**Figure 3:**
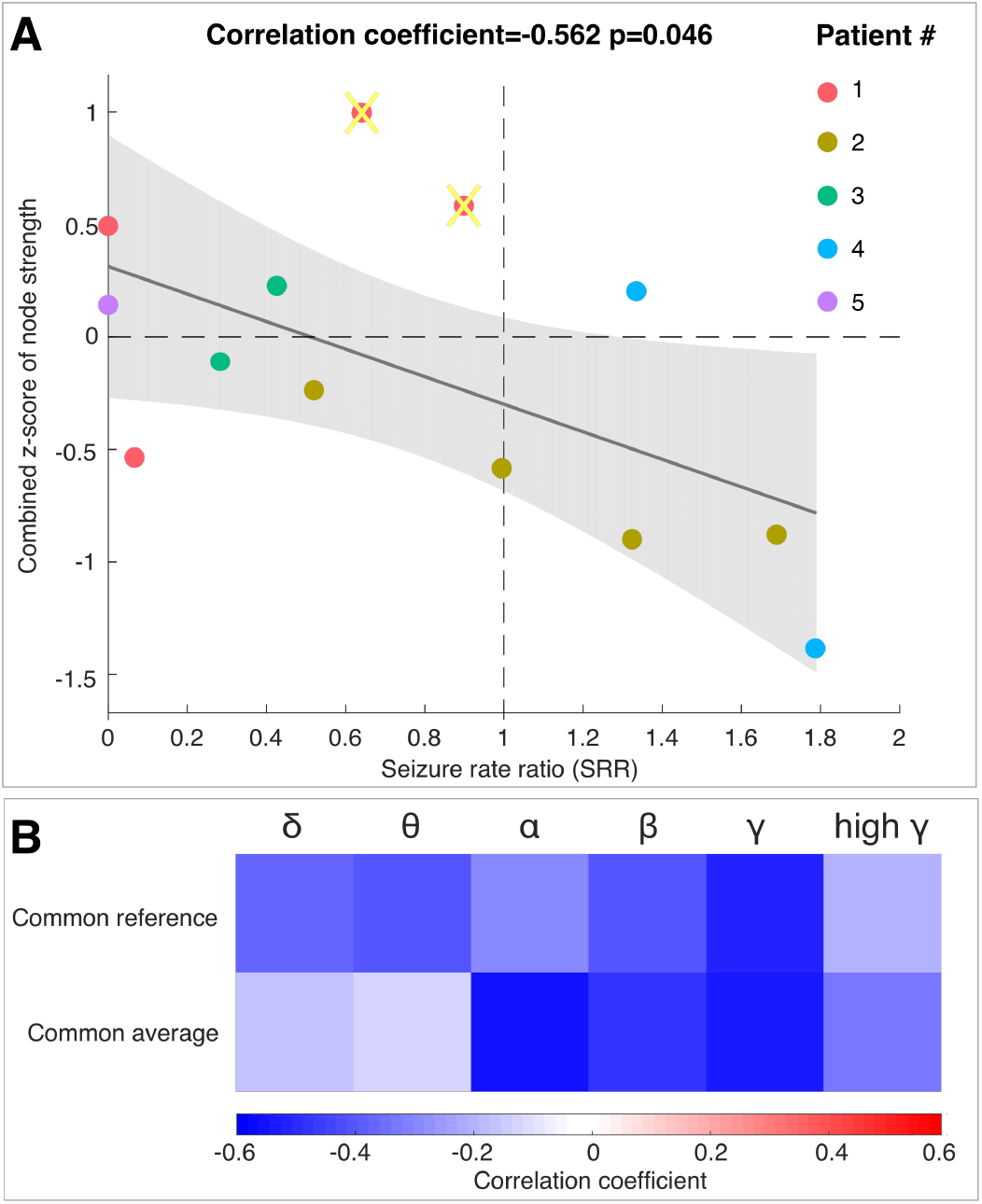
Relationship of baseline interictal network node strength and ensuing seizure rate ratio in SCS for all five patients. **A**: Scatterplot of baseline interictal network node strength (z-score) of the electrodes that were used for stimulation in SCS vs. the ensuing seizure rate ratio in the corresponding SCS. Solid grey line shows the regression, and grey shaded area shows the confidence interval of the regression. Two datapoints marked with a yellow cross indicate where the SCS condition was using 60 Hz as stimulation frequency, all other datapoints were 130 Hz stimulation. When excluding these two datapoints, the correlation coefficient becomes r=-0.711, p=0.014 **B**: The correlation coefficient (i.e. slope of regression) between z-score of node strength and seizure rate ratio is shown for networks generated using different montages and different frequency bands.

As a second step, we tested if the correlation using the alpha band data was consistent also for other frequency bands of the interictal baseline network. We additionally tested the common reference montage against the common average montage to assess robustness of results. Similar to previous work [16], we found that the interictal networks are reasonably consistent, and give similar results across both frequency bands and montages (Fig. 3B). All correlation coefficients of z-score node strength vs. SRR are negative, indicating the same relationship we found for the alpha band common average setting.

## Discussion

In summary, we demonstrated in five patients with epilepsy that baseline interictal network properties could be used to predict the effect of Subacute Cortical Stimulation (SCS) conditions on seizure rates during the stimulation. This represents a significant step forward in terms of developing brain stimulation as a targeted treatment for epilepsy patients, as it provides a data-driven method of identifying stimulation targets subject-specifically, based on interictal data.

The study of functional networks of interictal electrographic data such as EEG have a long history [25–30], with a series of recent publications pointing towards its clinical value (e.g. in localising the seizure focus) [14–16, 31]. The exact interpretation of the functional network depends on the study, the bivariate measures used, and the subsequent analysis applied to it. The measure and analysis we chose (correlation, and node strength) has a straight-forward interpretation: in Fig. 2 red colour means more correlation to other channels than average, and blue means less correlation (in the alpha band using a common average montage). Our results across five patients (Fig. 3) appear to suggest that stimulation delivered through red (high average correlation) electrodes in SCS is successful in diminishing number of seizures, while stimulation delivery through blue (low average correlation) electrodes may be associated with increments in seizure numbers.

One way these observations can be interpreted is that if we assume that SCS stimulation has an overall inhibitory/disruptive effect on the targeted cortical region, as suggested in [32–34], then disruption of areas with high average correlation (or high connectivity) to other regions can suppress seizures. This is also in line with our recent observation in a separate dataset that surgical removal of such high correlation regions appears to lead to seizure freedom [16]. Conversely, the disruption of areas with low average correlation has the opposite effect, which could be interpreted as these areas exerting seizure control that is then disturbed by the SCS.

Many studies have indicated clinical usefulness for the interictal correlation networks, but their exact physiological underpinning is still debated. One indication is that these networks are related to anatomical connections (particularly in the gamma band [35, 40]). In an animal model, it was further shown that early epileptogenesis is paralleled by an increase in network correlation interictally [36]. We hypothesise at this stage that there may be several cellular and circuitry mechanisms leading to abnormally high correlations interictally; however, these correlations themselves may serve as a dynamic and unifying hallmark of epileptogenicity.

It is also unknown how these interictal correlation networks are related to interictal spikes. We believe that it is unlikely that correlation networks are a direct result of co-occurrence of interictal spikes (also see [37, 38] for further support). The networks are relatively stable across all frequency bands [16] (especially theta, alpha, beta, and gamma), while interictal spikes are usually limited to specific bands. Nevertheless, the exact relationship between interictal spikes and interictal correlation networks should be elucidated in future studies to improve clinical and physiological interpretability.

The temporal variability of interictal correlation networks is also being investigated. In short-terms (around 100 seconds), it has been convincingly shown that these networks are stable [13, 25, 28, 30, 31], which is why we choose to average our networks across 500 seconds. However, in terms of variability over hours and days, very little work has been done, possibly due to the computational challenges. One study showed that there may be circadian and other rhythmic changes impacting these networks [39], while another study reports stability over days (using a different bivariate measure of time-lagged mutual information) [30]. Future work should investigate these temporal variabilities, especially in the context of seizure activity and their spatial relationship to the seizure onset and irritative zones.

Further technical limitations to be considered in our work include the interaction of SCS with anti-epileptic drugs, the relatively short duration of the SCS and hence difficulties in estimating accurate seizure rates, as well as small subject numbers. Despite these limitation, we believe that we have contributed an important new proof-of-principle observation: we report that in our cohort, interacting with the interictal network via SCS can augment or reduce seizure rates in a predictable manner based on the interictal network structure.

## Conclusions

We have demonstrated that interictal functional network properties contain information that can correlates with the outcome of neuromodulation in epilepsy patients. Future work will focus on the relationship between interictal functional networks and other clinical variables and markers. We hope that our work will enable prospective application of these interictal functional networks in designing patient-specific neuromodulation protocols, perhaps even in a wide range of neurological disorders.

## Abbreviations

SCS: subacute cortical stimulation
iEEG: Intracranial electroencephalography

## Acknowledgements

We thank Andrew Jackson, and Gabrielle M Schroeder, Roger Whittaker, Michael Mackay, and Dennis Prangle for fruitful discussions and feedback on the manuscript.

## Appendix

## Appendix A Consistency of network node strength over time

To analyse the temporal variability of our network node strength, we calculated the correlation networks on each 2 second time window, for each 500 second recording, as described in the Methods. To evaluate the variability within the 500 second recording, we averaged every 50 seconds of the recording, thus obtaining ten networks for which we calculated the node strength. These network node strengths were then z-scored across all channels. The average and standard deviation of the node strength z-score is shown in the subsequent figures A1-A5 as errorbar plots. The different 500 second recordings for each subject are shown as different colours in these plots. For all subjects, we can see that the node strength z-scores from different 500 second recordings agree well with each other in the same patient, although they are at least 4h apart from each other. Most channels are also within one standard deviation of the within-recording variability.

**Figure A1:**
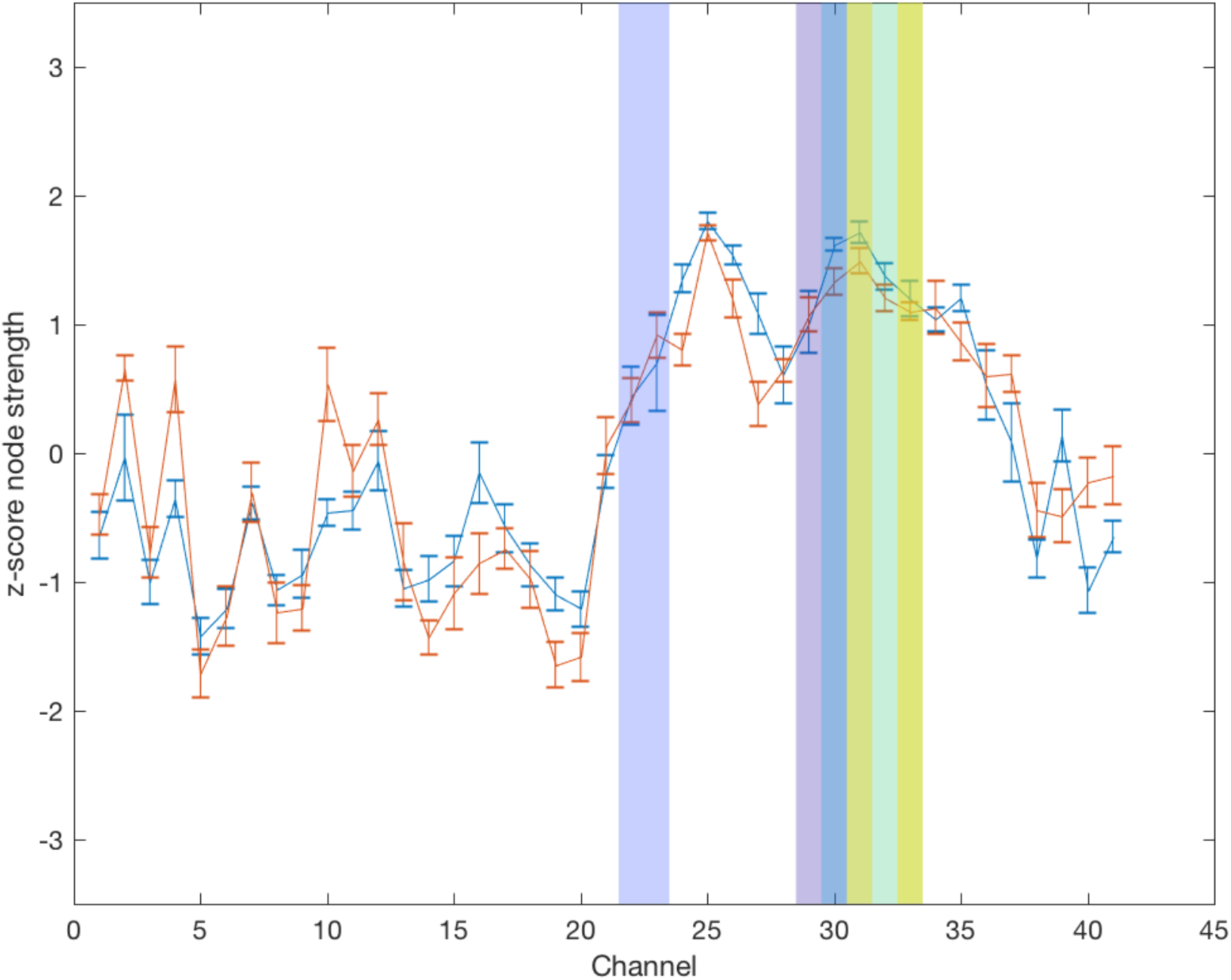
Patient 1. Errorbars of the same colour represent standard deviation in node strength z-score with the same 500 s recording. Different colours represent measurements from different 500 s recordings, at least 4 h apart. Vertical transparent stripes indicate different channels that were used for stimulation (each pair is shown with the same colour).

**Figure A2.**
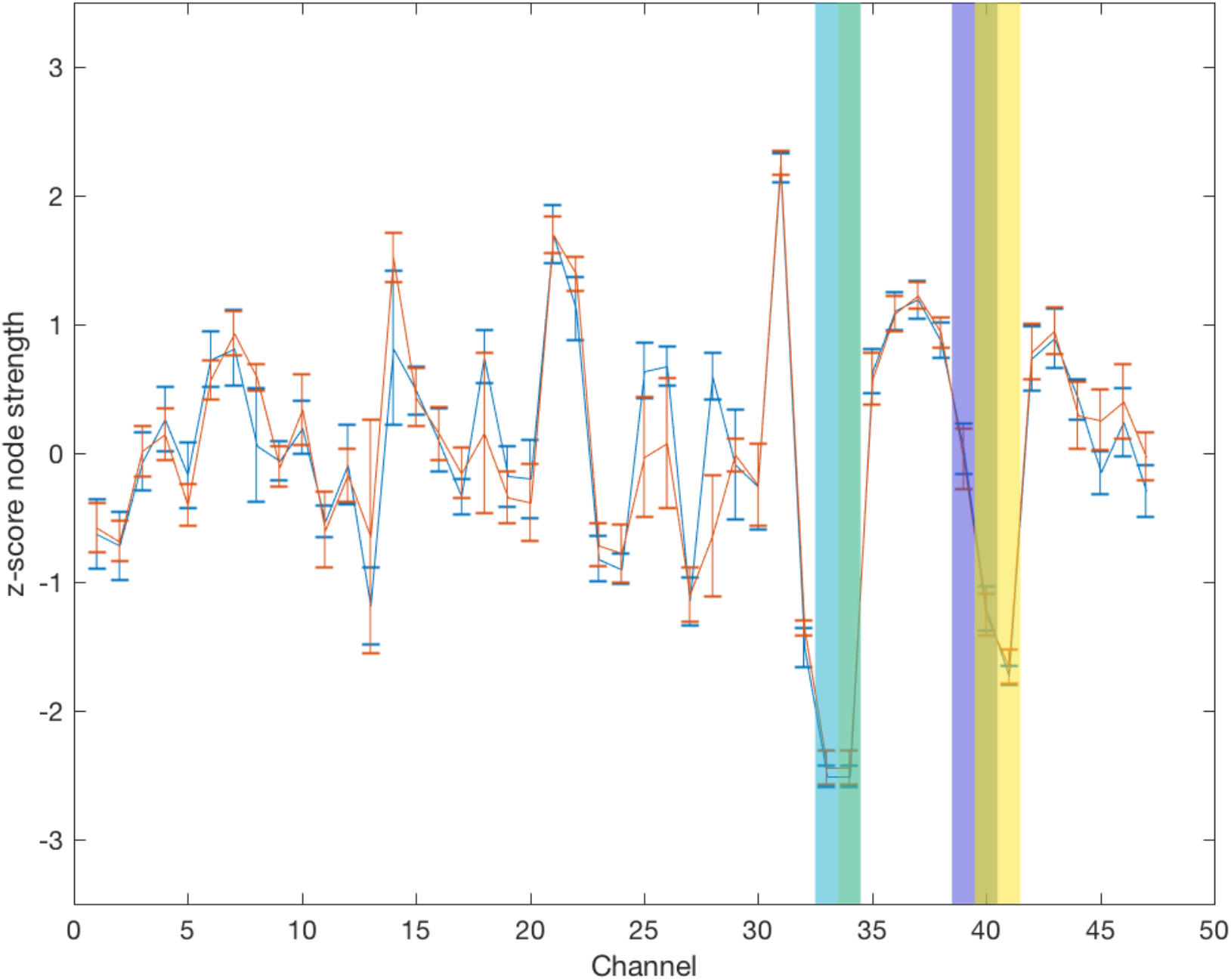
Patient 2. All other descriptions are the same as Fig. A1.

**Figure A3.**
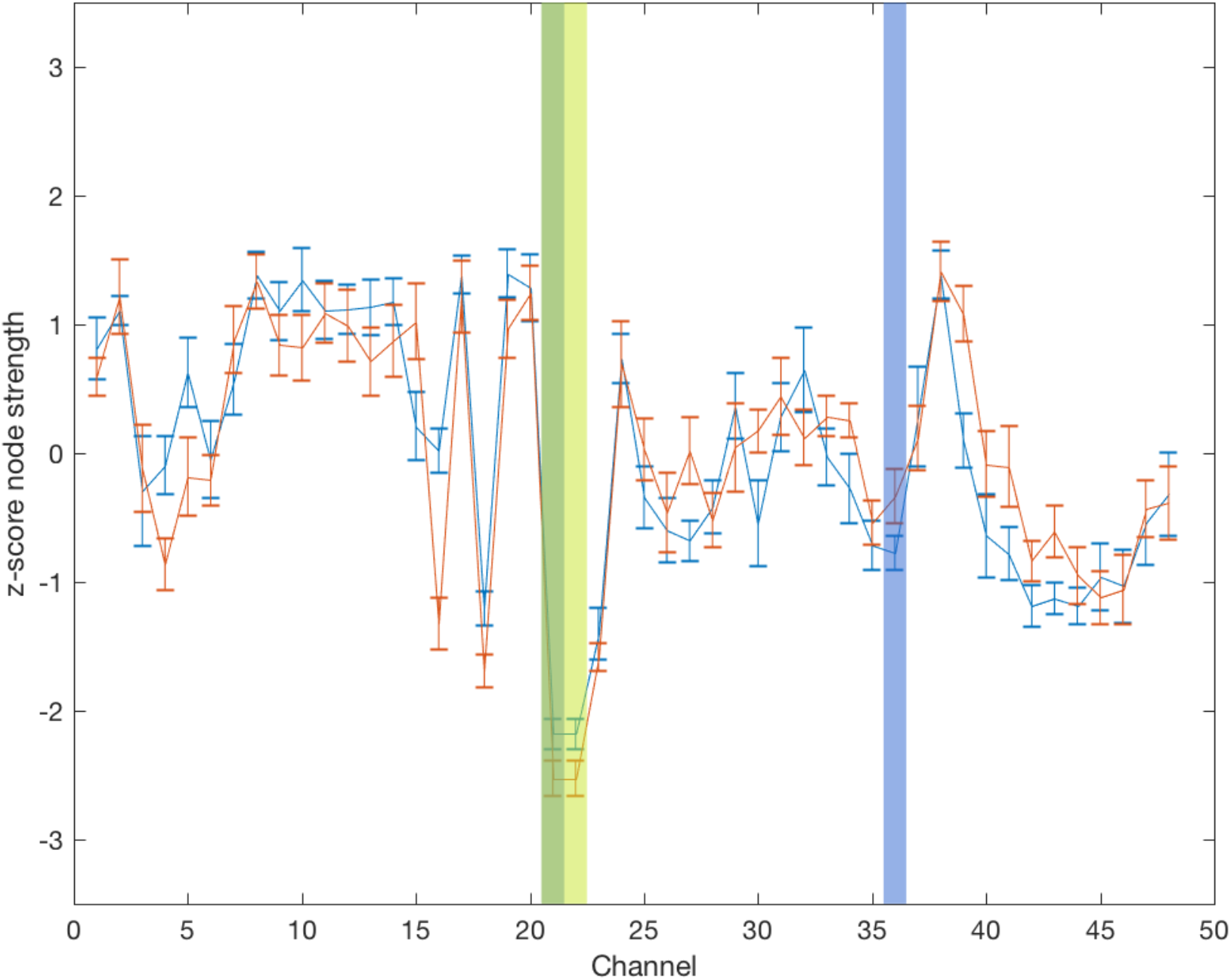
Patient 3. All other descriptions are the same as Fig. A1.

**Figure A4.**
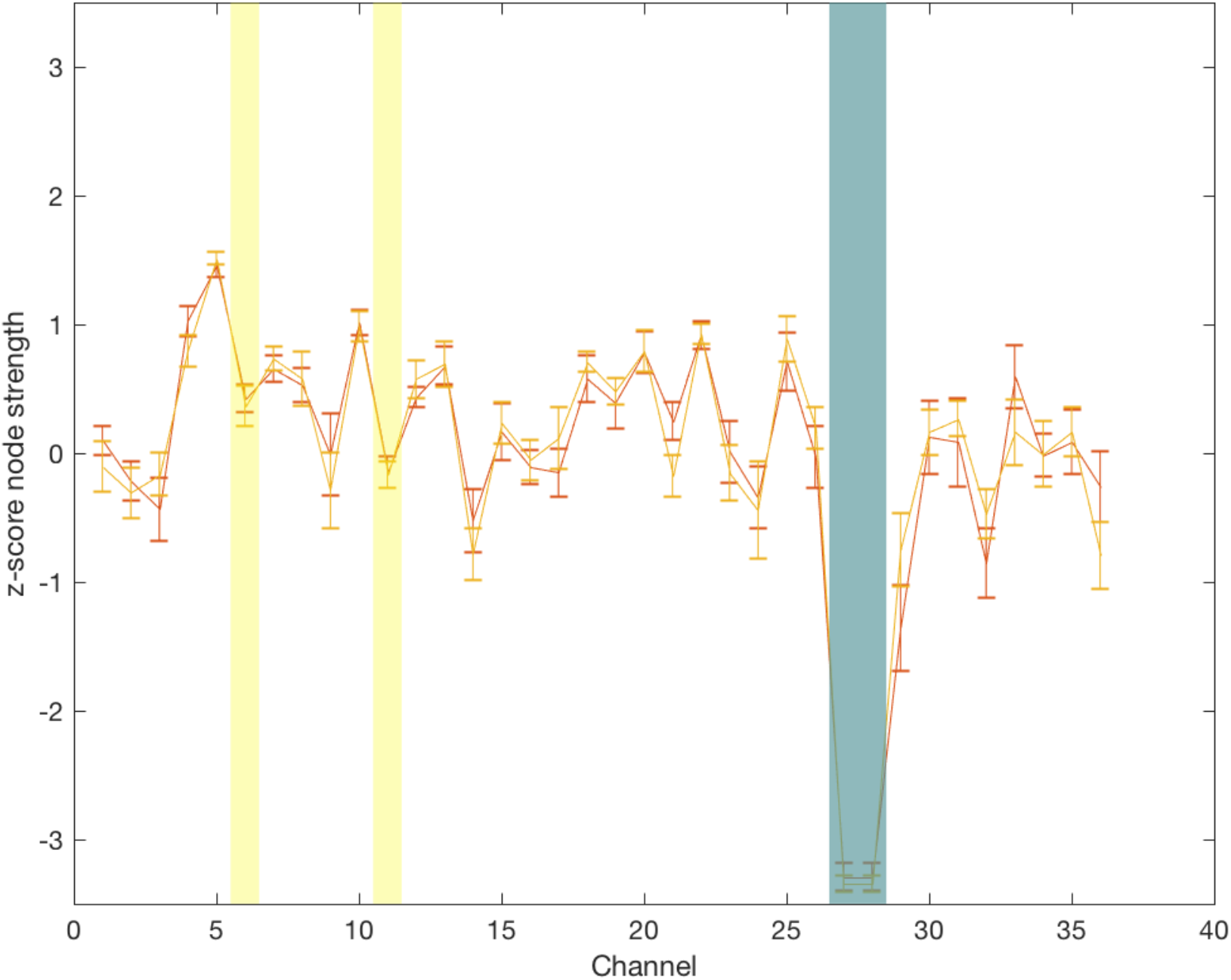
Patient 4. All other descriptions are the same as Fig. A1.

**Figure A5.**
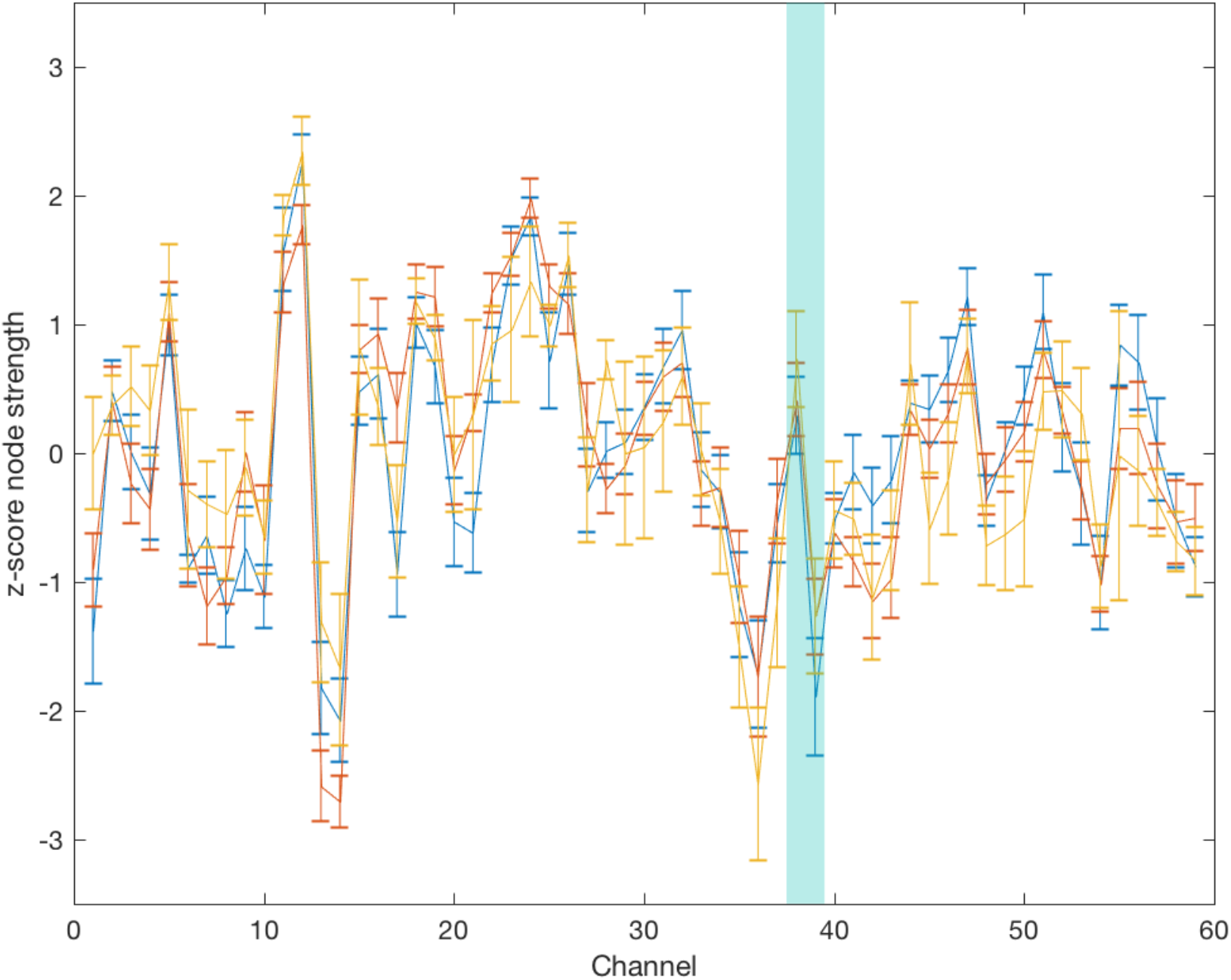
Patient 5. All other descriptions are the same as Fig. A1.

